# Optimized Monophasic Pulses with Equivalent Electric Field for Rapid-Rate Transcranial Magnetic Stimulation

**DOI:** 10.1101/2022.08.29.503248

**Authors:** Boshuo Wang, Jinshui Zhang, Zhongxi Li, Warren M. Grill, Angel V. Peterchev, Stefan M. Goetz

## Abstract

**Objective:** Transcranial magnetic stimulation (TMS) with monophasic pulses achieves greater changes in neuronal excitability but requires higher energy and generates more coil heating than TMS with biphasic pulses, and this limits the use of monophasic pulses in rapid-rate protocols. We sought to design a stimulation waveform that retains the characteristics of monophasic TMS but significantly reduces coil heating, thereby enabling higher pulse rates and increased neuromodulation effectiveness.

**Approach:** A two-step optimization method was developed that uses the temporal relationship between the electric field (E-field) and coil current waveforms. The model-free optimization step reduced the ohmic losses of the coil current and constrained the error of the E-field waveform compared to a template monophasic pulse, with pulse duration as a second constraint. The second, amplitude adjustment step scaled the candidate waveforms based on simulated neural activation to account for differences in stimulation thresholds. The optimized waveforms were implemented to validate the changes in coil heating.

**Main results:** Depending on the pulse duration and E-field matching constraints, the optimized waveforms produced 12% to 75% less heating than the original monophasic pulse. The reduction in coil heating was robust across a range of neural models. The changes in the measured ohmic losses of the optimized pulses compared to the original pulse agreed with numeric predictions.

**Significance:** The first step of the optimization approach was independent of any potentially inaccurate or incorrect model and exhibited robust performance by avoiding the highly nonlinear behavior of neural responses, whereas neural simulations were only run once for amplitude scaling in the second step. This significantly reduced computational cost compared to iterative methods using large populations of candidate solutions and more importantly reduced the sensitivity to the choice of neural model. The reduced coil heating and power losses of the optimized pulses can enable rapid-rate monophasic TMS protocols.

## 1. Introduction

Magnetic stimulation is a noninvasive technique to activate neurons [1]. Transcranial magnetic stimulation (TMS) is used to probe and modulate brain function [2,3], and clinical TMS routinely serves for the diagnosis and treatment of psychiatric and neurological disorders [4,5]. TMS uses an electromagnetic coil to generate an intense time-varying magnetic field of about one tesla to induce an electric field (E-field) in the brain [6]. Various TMS waveforms and pulse directions activate different neural targets and achieve different levels of changes in excitability [7–11]. For instance, repetitive TMS with monophasic pulses typically generates stronger facilitation and activates cortical neurons with higher directional selectivity [12–16], but clinical applications almost exclusively use biphasic pulses.

The reasons for exclusive use of biphasic pulses in repetitive TMS are mostly technical. Due to the weak electromagnetic coupling, TMS requires currents and voltages of several kiloamperes and kilovolts [17–20], which generate rapid coil heating during repetitive pulse trains. As safety regulations limit the temperature of the coil surface in contact with the subject [21], most coils for repetitive stimulation require complex cooling systems but still have heating-related restrictions on the pulse strength, rate, and/or number. As the least energy-efficient pulse shape, monophasic pulses cannot operate at high repetition rates due to the excessive power loss. This power loss causes heating in the coil and the hardware needs a sufficient power supply to provide this power. Biphasic pulses, despite producing weaker facilitation and being less selective, can be generated more efficiently, cause less heating, and therefore allow higher repetition rates [22,23].

For example, quadripulse stimulation (QPS) with four monophasic pulses delivered in a rapid sequence [24] creates the largest [25–27] and most repeatable [28,29] changes in excitability, but is typically performed with low burst repetition rate such as 0.2 Hz [30,31]. QPS using higher burst rate (e.g., ∼ 5 Hz) appears to retain its efficacy with just a fraction of the stimulation session duration [32,33]; however, such an accelerated protocol has only been demonstrated with the less effective but more energy efficient biphasic pulses. Similarly, theta-burst stimulation with monophasic pulses is currently not possible. Present commercial devices cannot generate any of the approved treatment protocols for major depression, which require pulse rates of at least 10 Hz [34–36], with monophasic pulses, and all medical treatments use biphasic pulses despite the resulting smaller changes in excitability. Repetitive TMS could be significantly improved with a pulse shape that produces changes in excitability equivalent to those produced by a monophasic pulse and but with lower loss and higher energy efficiency.

The high power requirements of TMS devices and associated extreme demands on the underlying electronic circuits and passive components are a key reason for the slow technological development of pulse sources that allow control of the pulse waveform [22]. Therefore, minimization of the energy required to generate specific E-field waveforms is advantageous. Previous theoretical optimization with nonlinear neuron models suggested that the pulse energy and coil heating could be reduced by more than a factor of two [37]. However, the reported pulses are more bidirectional than conventional monophasic pulses and bear no close resemblance with any conventional or clinically approved pulse shape, and thus it is not clear that they could produce equivalent changes in excitability.

Several TMS technologies allow increased waveform flexibility [38–45], but pulse shapes that are both highly efficient with low coil heating and replicate accurately the E-field waveform of monophasic pulses have not been implemented. Herein we design pulse shapes that produce estimated neural activation equivalent to conventional monophasic pulses by matching the induced E-field but with minimized pulse power and coil heating. We introduce a novel model-free optimization approach, and nonlinear models are introduced only for amplitude adjustment, thereby reducing substantially the computational demands and sensitivity to model selection.

## 2. Method

### 2.1. Optimization Rationale and Overview

Unlike electrical stimulation, the temporal waveform of the TMS E-field, *E*(*t*), and current density induced in the tissue are not directly proportional to the output current generated by the TMS device, *I*_coil_ (*t*), but rather proportional to its time derivative [46]:

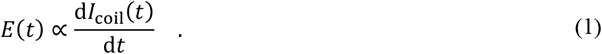

Therefore, a slow shift in the baseline of the coil current *I*_coil_ before the pulse minimally affects the induced E-field but reduces the energy loss, *W*, [37], which is mostly in the form of ohmic heating of the coil, its cable, and other circuit components in series with the coil per

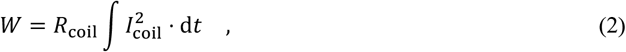

where *R*_coil_ is the total resistance dominated by the coil. Specifically, for monophasic TMS pulses with unidirectional *I*_coil_, shifting the entire waveform reduces the peak amplitude of *I*_coil_ and *W*. Through a tightly enforced shape similarity of the resultant E-field, the optimized pulse should have similar stimulation effects compared to the monophasic TMS pulse (Figure 1A), and preserve stimulation characteristics of the latter, such as directional selectivity and motor activation latency [9,47].

**Figure 1:**
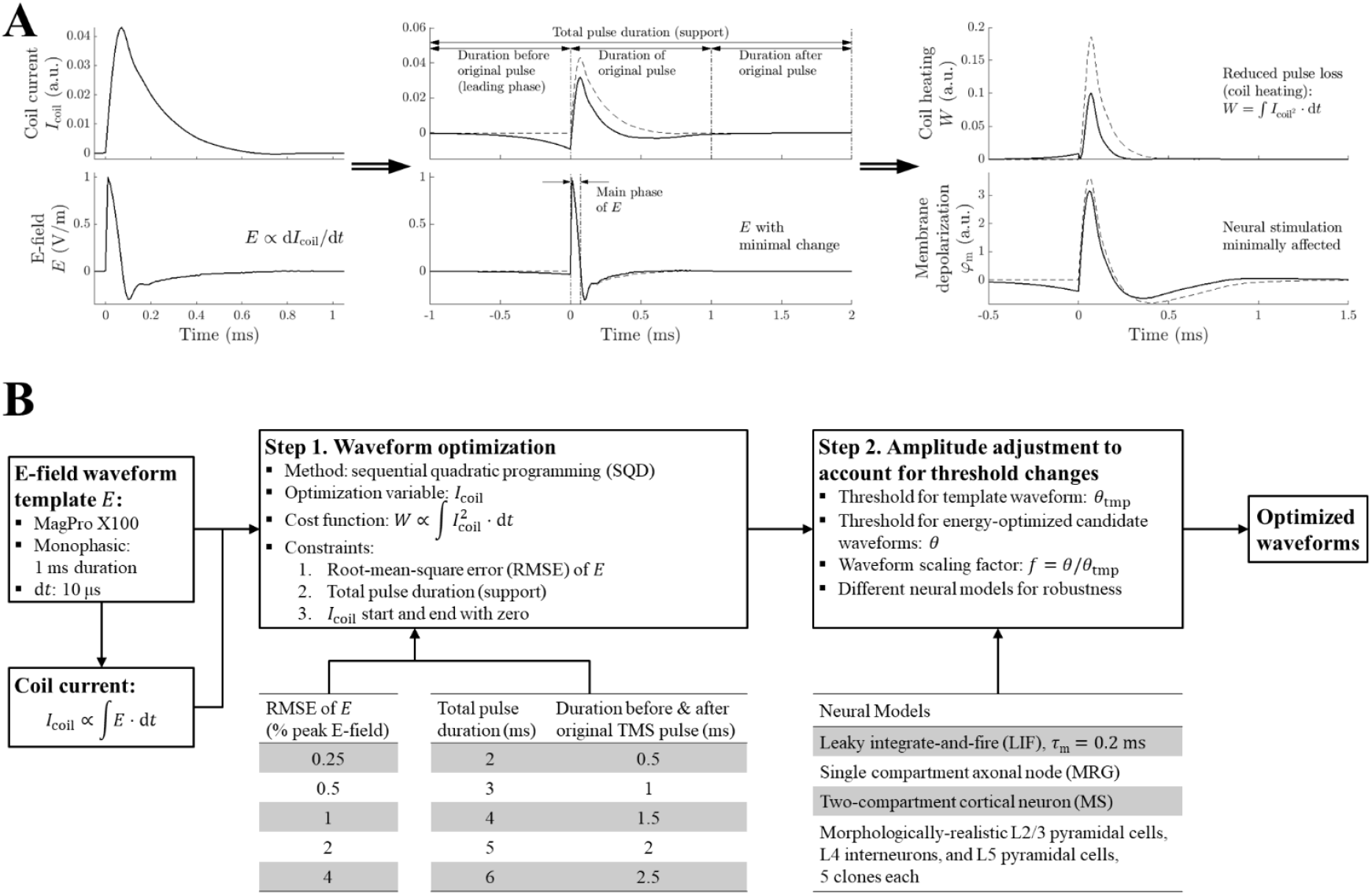
Waveform optimization principles and steps. **A**. The waveform optimization used the relationship between the TMS-induced E-field waveform *E* and coil current *I*_coil_. A shifted baseline reduced the amplitude of the current waveform and ohmic losses, whereas the E-field waveform and its stimulation effect were only minimally affected. **B**. The optimization started with an E-field waveform template, and, in the first step, produced a corresponding current waveform with minimal heating while limiting the error of the E-field compared to the template. The total pulse duration represents the length of the compact support, i.e., the time interval in which the optimization can let the current can be non-zero. In the second step, the amplitude of the candidate waveforms was scaled to account for differences in neural activation thresholds introduced during the first step. a.u. arbitrary units. MRG: McIntyre–Richardson–Grill model. MS: Mainen– Sejnowski model.

We set up a free optimization, where loss-reducing features of the current waveform can evolve just through the objective and the constraints without introducing any *a priori* features. The optimization problem is separated into two steps. The first step is formulated as a classic optimization problem that focuses on modification of the waveform shape; the second step deploys neural models to adjust the waveform amplitude to account for minor shifts of the activation threshold that might arise from the first step (Figure 1B).

### 2.2. Two-Step Optimization

The optimization problem started with the *I*_coil_ and the *E* of the conventional TMS monophasic pulse shape. We used an E-field waveform recorded from a MagPro X100 device (MagVenture A/S, Farum, Denmark), which was normalized to one (V/m) at the pulse start, and then calculated the corresponding coil current (in arbitrary units) as the integral (cumulative sum) of the E-field waveform. Half-sine and biphasic pulses of the same TMS device were also used for comparison. The optimization algorithm used a sequential quadratic programming method, implemented with the MATLAB function `fmincon`, and worked on *I*_coil_ to minimize the cost function *W* (with *R*_coil_ set to unity). Either the original monophasic pulse or random Gaussian noise was used to initialize *I*_coil_. To ensure minimal effect on the E-field and subsequent neural activation, the main optimization constraint was the root-mean-square error (RMSE) of the E-field waveform compared to the original TMS template waveform. We performed optimization with five RMSE levels: 0.25%, 0.5%, 1%, 2%, and 4% of the peak E-field amplitude. Constraints were also imposed at the beginning and the end of the pulse, so that *I*_coil_ started from and returned to zero. Thus, the pulse current had a mathematically compact support, which ensured a finite pulse duration and avoided any unaccounted pulse energy or loss. For computational optimization purposes, the waveforms were sampled with a 10 μs time step. This temporal sampling turned out to be sufficient as doubling the sampling (5 μs time step) did not affect the optimization (Figure S1).

Another constraint in the optimization was the duration of the optimized pulse, which—together with the RMSE constraint—determined how long before and after the reference pulse additional features can evolve. For example, a low-amplitude, slowly changing leading phase that shifted the current baseline before the depolarizing main phase emerged in our prior work on pulse optimization without shape constraints [37]. The original TMS current pulse had a duration of approximately 0.75 ms due to the long current decay, whereas the dominant phase of the E-field was only about 75 μs long (Figure 1A). We extended (zero-padded) the original TMS pulse (set to 1 ms total duration) both before and after with five durations between 0.5 ms to 2.5 ms, resulting in a total pulse duration between 2–6 ms. These durations represent the length of the compact support, i.e., the maximum interval in which the optimization can shape a pulse, and corresponded to 201 to 601 time points for *I*_coil_ and *E*. A desktop computer (with an Intel Core i7-8700 CPU and 16 GB RAM) performed each optimization condition independently up to ten times to examine the consistency of the results and the runtime. Computation and data analysis were performed with MATLAB (version 2021a, The Mathworks, Natick, MA, USA).

The second step in the design process scaled the amplitude of the candidate waveforms to the stimulation threshold in neural models. We delivered the E-field waveforms of the original TMS template and the candidate waveforms to neuron models and calculated their respective stimulation thresholds *θ*_tmp_ and *θ* within 0.1% accuracy using a binary search. To allow sufficient time for the stimulation to evoke an action potential, the simulations were extended by 6.5 to 8.5 ms depending on the total duration of the pulse. For accurate simulation of thresholds, the stimulation waveforms were linearly interpolated with a time step of 2.5 μs. We scaled the amplitude of each candidate waveform by its corresponding factor

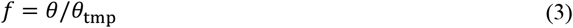

to form the optimized threshold-matched waveforms. We quantified the change in loss of the threshold-matched pulses as a percentage of the original monophasic pulse

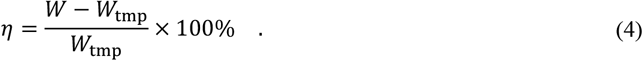

### 2.3. Neuron models

Several neural models were used to assess the robustness of the second, amplitude adjustment step. The first model was a single-compartment linear leaky integrate-and-fire (LIF) model. A time constant of 200 μs was chosen based on experimental neural responses to TMS [44,48,49]. The second was a single compartment model (MRG, McIntyre–Richardson–Grill) of a peripheral axon node [37,50]. The third was a two-compartment model (MS, Mainen–Sejnowski) that replicated firing pattern of cortical neurons with realistic dendrite morphology [51,52]. An intracellular current injection proportional to the E-field waveform stimulated these models, which were simulated in MATLAB on the same desktop computer that performed the optimization.

The remaining models were 15 morphologically-realistic cortical neuron models from the Blue Brain project [53] adapted for studying brain stimulation in the human cortex [47,54], including layer 2/3 pyramidal cells, layer 4 interneurons, and layer 5 pyramidal neurons, with five morphological clones used for each layer. The simulations were set up in MATLAB and performed in NEURON (version 7.7) [55] on the Duke Compute Cluster. The extracellular stimulation was applied using quasi-uniform approximation of the E-field [56]. E-field orientations were sampled in a spherical coordinate system in which the *z*-axis aligned with the somatodendritic axis [47,57], with 13 polar angles in [0°, 180°] (average spacing of 15°), 18 azimuthal angles in [0°, 360°) (spacing of 20°), and a total of 200 (11×18+2) orientations. For each cell model, amplitude adjustment factors were calculated for all 5000 (200×5×5) combinations of orientations and candidate waveforms.

### 2.4. Experimental waveform implementation

A modular pulse synthesizer TMS (MPS-TMS) device with six modules [41] was used to generate the optimized pulses. All waveforms were normalized to the maximum peak value and interpolated using a 5 MHz sampling rate. Before each pulse delivery, the internal capacitors of the modules (3.986 mF per module) were charged to 75 V, which generated output voltage between −450 V and 450 V. Despite the capacitors being disconnected from the dc power supply during the pulse, the control algorithm compensated for the voltage drop to maintain the correct amplitude of the current delivered.

A custom minimum-energy figure-of-8 coil following a previous design [58] served as the test coil for the pulses. The coil had an inductance *L*_coil_ of 9.46 μH and resistance *R*_coil_ of 11.7 mΩ at 1 kHz, measured in four-wire configuration with an LCR/ESR meter (Model 889A, B&K Precision Corporation, USA). Electrical measurements were performed at 100 MHz sampling frequency with an oscilloscope (Tektronix MDO3054, Beaverton, OR, USA). A Rogowski current transducer (CWT 60B, Power Electronic Measurement Ltd., Nottingham, UK) and high-voltage differential probes (Tektronix THDP0100) measured the coil current and voltage, respectively. The module voltage *V*_DC_ was also measured from the modules’ dc capacitors after each pulse. A printed-circuit-board-based triangular probe of 70 mm height [59,60] measured the E-field corresponding to a depth of 15 mm under the center of the coil in an 85 mm sphere approximating the head. For visualization only, a 3^rd^-order Butterworth low-pass filter with a cut-off frequency of 100 kHz was used to filter the recorded waveforms (with MATLAB function `filtfilt`).

The losses for all pulses were calculated from the measurements with three similar metrics: the integral of the coil current squared multiplied by the coil resistance obtained from the LCR/ESR meter measurements (2), the energy dissipation based on instantaneous power

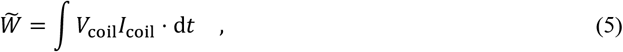

and the change in the stored energy using the pre- and post-pulse voltage of the device’s dc capacitors

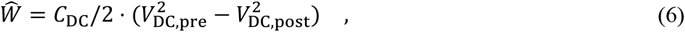

where the total capacitance of all six modules *C*_DC_ was 23.92 mF and *V*_DC,pre_ was 75 V. The first metric captured mostly ohmic heating of the coil; the second included also other losses, such as mechanical vibration of the coil and electromagnetic field radiation; and the third incorporated further operational losses internal to the MPS-TMS system. The pulse energy, which was the minimum energy required to generate the pulses in the TMS coil, was also calculated using

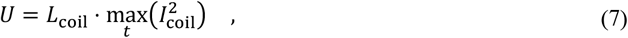

which corresponded to the peak magnetic field energy in the TMS coil, or

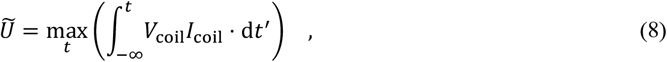

which included both the reactive energy *U* that was provided to the coil and recovered by the device at the end of the pulse and the active power dissipated up to the time of the magnetic field reaching its peak strength.

## 3. Results

For clarity, the figures in the main text show a subset of the results, e.g., three of the five RMSE constraints, two or three of the five duration constraints, and one or two model clones for each cell type of the Blue Brain models. The full datasets are provided in the supplement.

### 3.1. Waveform optimization

The optimization generated current waveforms with a negative leading phase that shifted the onset of the main phase toward a more negative baseline. The peak amplitude was decreased and the shape was narrower compared to the original monophasic pulse (Figures 2A and S2A). The magnitude of the leading phase increased and the pulse width and amplitude of the main phase decreased, respectively, with larger RMSE constraint on the E-field waveform and longer pulse duration. The optimized E-field waveform consisted of a main phase that approximated the original E-field waveform but had a slightly reduced peak amplitude and a negative leading phase with amplitude between −0.4% and −35.4% of the corresponding main peak (Figures 2A and S2A). The changes in loss compared to the original TMS pulse ranged from −15% to −95% (Figures 2B and S2B, dashed lines in bottom left panel). Initializing the optimization with random noise resulted in the same optimal waveforms (within the optimization tolerance) as initializing with the monophasic TMS pulse.

**Figure 2:**
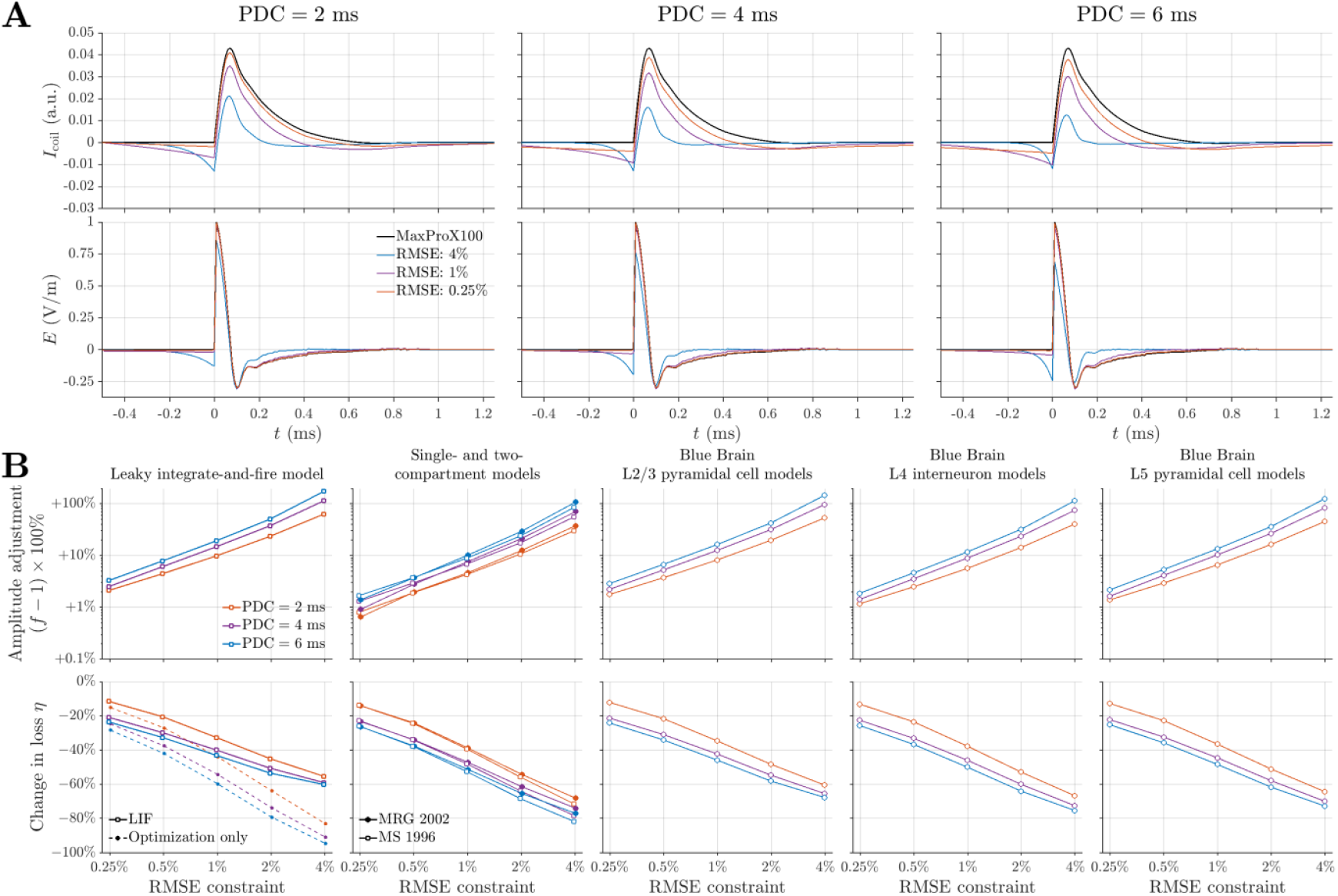
Waveform optimization and amplitude adjustment steps. **A**. The optimized coil current waveforms (colored lines) had a negative leading phase, reduced amplitude, and narrower shape than the original TMS monophasic pulse (black lines). The corresponding E-field waveforms included a negative leading phase but their main phase resembled the E-field produced by the original pulse. Only the center portion of the optimized waveforms are shown, time-shifted to match the monophasic pulse. **B**. The scaling factors *f* for adjusting the amplitude of the optimized waveforms to reach activation threshold are converted to percentage increase (*f* − 1) × 100% and shown on logarithmic scale. They increased with both constraints on the total pulse duration and the E-field RMSE, whereas the loss compared to the original TMS pulse decreased with these parameters. The loss without the amplitude adjustment are also shown in the first column (dashed lines). The results for the Blue Brain models are given for the first model clone for each cell type and are the mean across all the 200 E-field orientations. Results for all parameters and model clones are given in Figure S2. a.u.: arbitrary units. PDC: pulse duration constraint. LIF: leaky integrate-and-fire model. MRG: McIntyre–Richardson–Grill model. MS: Mainen–Sejnowski model.

The second step in the optimization adjusted the amplitude of the optimal waveforms based on the threshold to activate a range of neural models (Figures 2B and S2B). The threshold-based scaling increased the amplitude by a factor *f* of ×1.006 for the smallest RMSE and shortest pulse duration constraints and up to ×2.71 for the largest RMSE and longest pulse duration constraint. For a given parameter combination of RMSE and pulse duration constraints, the LIF model required the largest adjustment to the pulse amplitude (×1.02 to ×2.71) and thus had the least reduction in heating (−12% to −60%) among all the models. In contrast, all the nonlinear models, regardless of complexity, required smaller amplitude adjustments (×1.006 to ×2.33) and the waveforms produced less coil heating (−12% to −81%), and these were also highly consistent across the different neuron models. The amplitude adjustment factors *f* differed by only −0.3% to −19% between the two-compartment MS model and the Blue Brain models, and by −0.6% to −8.6% between the single compartment MRG model and the Blue Brain models, with larger differences for larger RMSE and longer pulse duration constraints. The coefficients of variation (i.e., ratio of standard deviation to mean) of the amplitude adjustment *f* across all 200 E-field orientations for each individual Blue Brain model (clone) and across the results (15×200) of all 15 models were between 0.5% to 10%, with higher values for larger RMSE and longer pulse duration constraints. The consistency of threshold adjustment across neural models, and especially across E-field orientations for each Blue Brain model, demonstrated that the optimization indeed generated stimulation pulse shapes that have the same characteristics as monophasic TMS pulses, such as directional selectivity. The leading phase of the pulse introduced a delay in neural activation when the action potentials generated by the optimized pulses were compared to those by the conventional monophasic pulse, with the delay equal to the duration the leading phase (Figure S3). Considering the main phase only, the optimized pulses had on average a zero delay, with some small variability across different E-field orientations for the Blue Brain models.

The mean amplitude adjustment of the Blue Brain models for each parameter combination was used to scale optimized candidate waveforms to generate the final optimized waveforms (Figures 3 (dashed lines in middle and bottom rows) and S2C). With their final RMSE only slightly increased from the corresponding RMSE constraints (Figure S4A, on average 0.27%±0.01%, 0.55%±0.01%, 1.13%±0.01%, 2.38%+0.05%, and 5.30%±0.26%, respectively, for 0.25%, 0.5%, 1%, 2%, and 4% RMSE constraint), these waveforms reduced coil heating by 12.5% to 75%. Some of the optimized monophasic pulses exhibited lower losses than conventional biphasic pulses, which already have lower stimulation thresholds [9,47] and less heating compared to conventional monophasic pulses (Figures 4A and S5). For this comparison, we used the biphasic pulses with reversed current direction compared to the monophasic pulse (Figure 4B), which are known to have a lower threshold due to the dominant second phase of the E-field [9,61,62].

**Figure 3:**
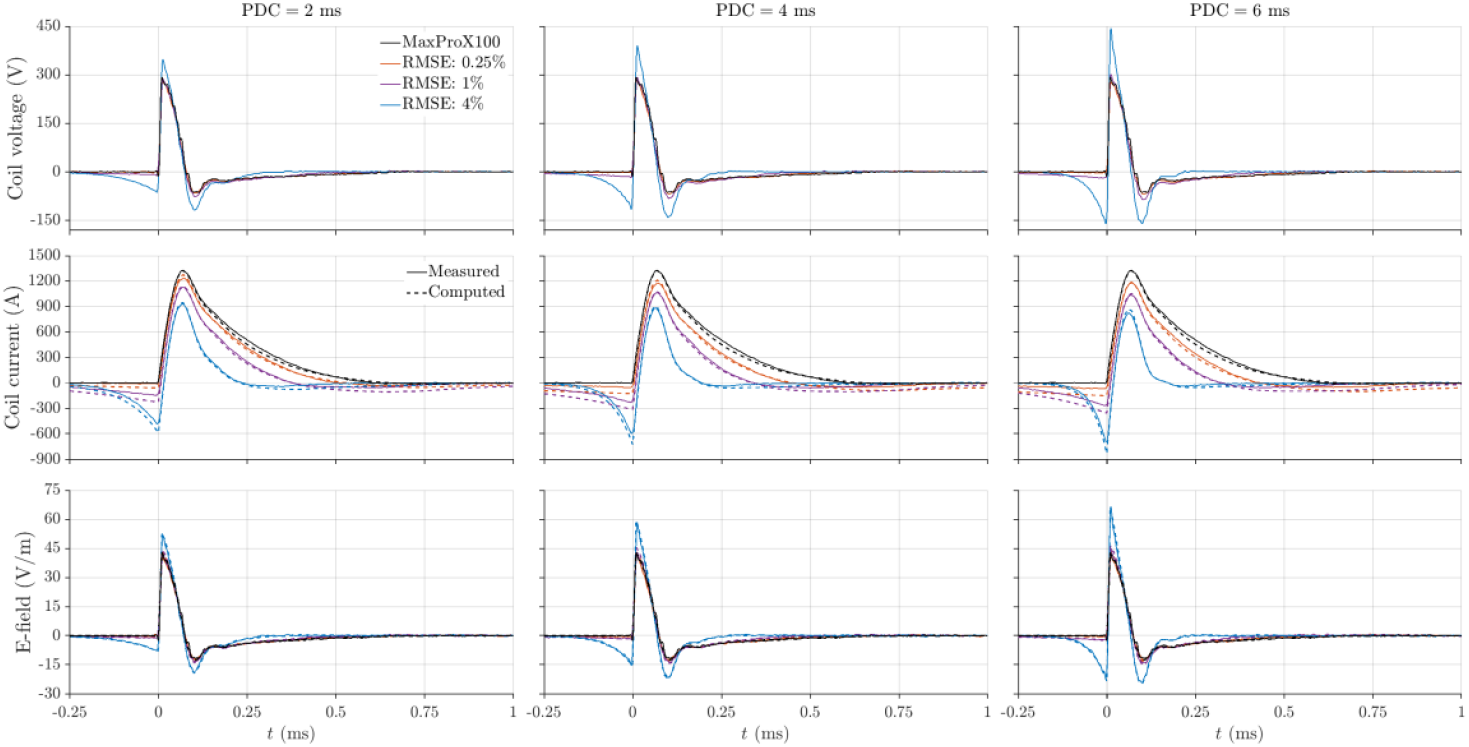
Computed and measured optimized TMS waveforms. The coil current and E-field waveforms show good agreement between the measurement (solid lines) and the computed results (dashed lines), which were scaled to match the peak of their measured counterparts (results without scaling shown in Figure S2C). Measurements were low-pass filtered for visualization only. Results for all RMSE and pulse duration constraints (PDC), the unfiltered measurements, and the effect of filtering are shown in Figures S7, S8, and S9−S10, respectively.

**Figure 4:**
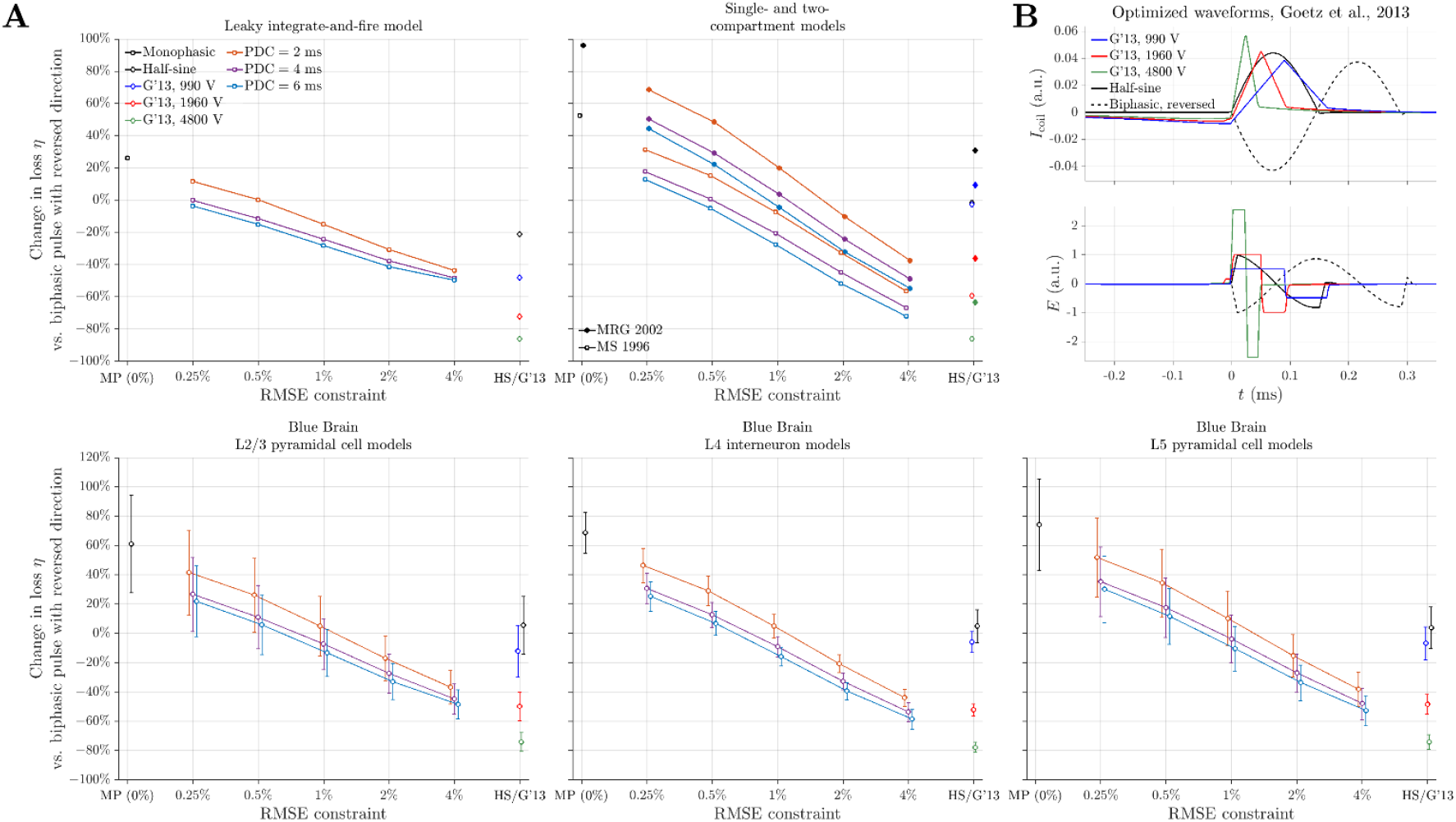
Difference in coil heating comparing pulses with monophasic, optimized, half-sine, and triangular shapes against biphasic pulse. **A**. The coil heating at stimulation threshold was calculated for a positive pulses for the monophasic (MP) and optimized pulses, as well as half-sine (HS) pulse and triangular optimal waveforms (G’13) from previous work [37], and for a biphasic pulse with reversed current direction (negative pulse). Similar format as bottom row of Figure 2B.. **B**. Three optimized waveforms from previous work [37], normalized to the second waveform (red line), and conventional half-sine and (reversed) biphasic pulses recorded from MaxPro X100 (black lines). The optimized waveforms have a total duration (support) of 1.6 ms with a 1.2 ms leading phase, with the voltage values indicating the optimization constraint on the coil voltage. Results for all pulse duration constraints (PDC) and model clones as well as full duration of optimized waveforms from previous work are shown in Figure S5.

### 3.2. Computational Runtime

The runtime of the E-field matching optimization step was between a few seconds and up to two minutes, with longer runtime for the optimization of longer pulse duration constraints (Figure S6, solid lines in leftmost column). Doubling and halving the sampling time step from 10 μs to 5 μs or 20 μs changed the optimization time by ×(10.7±3.3) (range ×6.0 to ×18.0) and ÷(7.4±3.0) (range ÷3.9 to ÷14.0), respectively (Figure S6, dashed and dotted lines in leftmost column). In comparison, the runtime of the threshold search varied for the different neural models, with single- and two-compartment models requiring 1–4 s and approximately 10 s, respectively (Figure S6, second column). The runtime of the Blue Brain models (Figure S6, third to fifth columns) depended on their morphological complexity, with the simpler L2/3 pyramidal neurons requiring less than 5 minutes, L5 pyramidal neurons typically completing within 10 minutes, and the most complex L4 interneuron requiring up to 30 minutes.

### 3.3. Waveform implementation

The measured waveforms (Figures 3 and S7, solid lines) closely matched the optimized waveforms (Figures 3 and S7, dashed lines), and the RMSE was between 0.8% and 4% for filtered waveforms and approximately 4% for unfiltered results (Figure S4B), indicating successful implementation of the optimized TMS pulses using the MPS-TMS device. The E-field peaked at approximately 66 V/m for the optimal waveform with highest amplitude (4% RMSE and 6 ms pulse duration constraint) and at 43 V/m for the original TMS pulse, with corresponding peak (filtered) coil voltages of 450 V and 292 V, respectively. The coil current, however, was lower for the optimized pulse (peak of 824 A versus 1323 A) due to the negative leading phase (negative peak of −690 A). The optimized waveforms generated less coil heating compared to the original TMS pulse (Figures 5A and S11A) and required less energy to generate (Figures 5B and S11B). For the various pulses, approximately two thirds of the loss 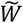 was in the form of heat *W* (66%±6%, range 51%−73%) and the energy efficiency of the MPS-TMS system 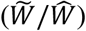 was 51%±2% (range 47%−55%). Despite some deviation, likely due to recording noise, the percentage difference of pulse energy *η* calculated from the measurement also matched the calculations for both the loss and total energy of the optimized waveforms (Figures 5C and S11C, and 5D and S11D, respectively).

**Figure 5:**
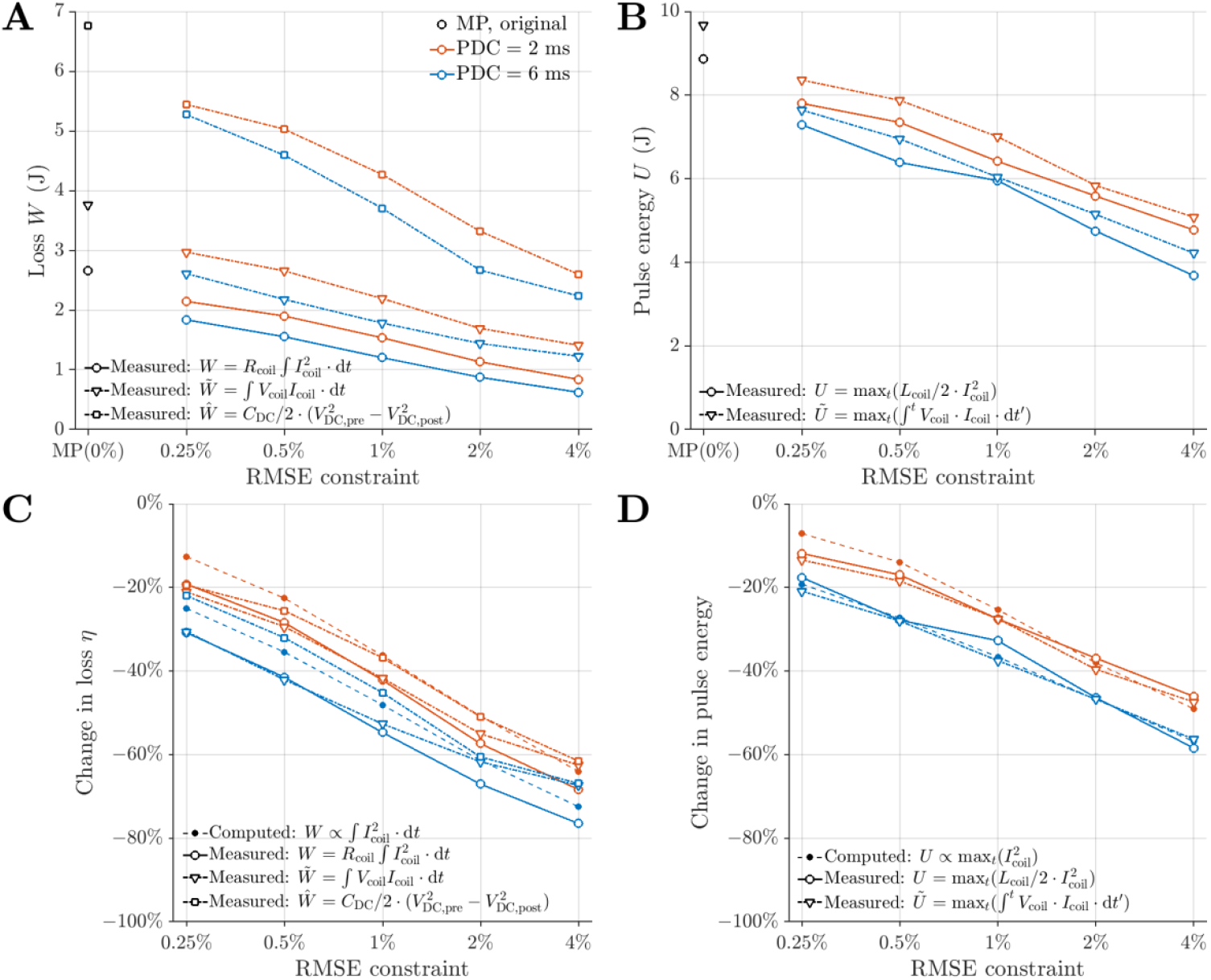
Losses and pulse energy of optimized pulses and original TMS pulse. **A**. The losses 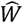 (square markers) included the internal losses of the MPS-TMS operation and the energy dissipated through the coil 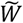 (triangle markers), which consisted of ohmic heating of the coil *W* (circle markers) and other losses. **B**. The total energy required to generate pulses. **C**. The losses of the optimized pulses compared to the original monophasic (MP) TMS pulse. **D**. The pulse energy of the optimized pulses compared to the original monophasic TMS pulse. Results for all pulse duration constraints (PDC) are given in Figure S11.

## 4. Discussion

### 4.1. Waveform optimization

We used a two-step approach to optimize the TMS pulse for minimal coil heating while retaining the excitation characteristics of conventional monophasic pulses. In the first step, a classic (i.e., non-heuristic) optimization problem was formulated and solved, and this resulted in optimal waveforms that exhibited a negative leading phase as predicted by the optimization rationale and observed in previous optimization studies [37,63]. The negative leading phase of both the current and E-field was not pre-defined in the optimization problem, but rather emerged as a result of the cost function and E-field constraint, as well as the increased pulse duration compared to the monophasic pulse template. As the problem was well-defined and deterministic, the optimization was robust and not subject to random factors such as parameter initialization.

The pulse shape optimization problem had two constraints: the RMSE of the E-field and the total pulse duration. The deviation of the E-field from the conventional monophasic pulse shape was limited to 0.25% to 4% RMSE during the pulse shape optimization and the final RMSE after the amplitude adjustment step was between 0.27% and 5.6% for modeling results and between 0.8% and 4.6% for measurements. Such errors are similar to typical pulse-to-pulse and machine-to-machine variability of common TMS devices [64] and are expected to produce comparable physiological effects. The RMSE constraint had a larger influence on the reduction of coil heating than did the pulse duration constraint (Figure 2B): relaxing the RMSE constraint (higher RMSE) reduced the heating more than increasing the non-zero pulse waveform duration. This was mostly due to higher RMSE resulting in a current waveform that had a main phase with significantly shorter duration and lower amplitude compared to the original TMS pulse, and shorter TMS pulses are known to require less energy and produce less heating [37,48,65,66]. For long pulse duration constraint, the further heat-reducing effect decreased, e.g., increasing pulse duration from 4 ms to 6 ms resulted in a smaller change of coil heating than increasing the pulse duration from 2 ms to 4 ms.

The short delay in neural activation introduced by the leading phase and longer pulse duration is not a problem in most TMS applications. When the optimized waveforms are used in place of monophasic pulses in repetitive TMS protocols, most of the leading phase and the prolonged tail of the pulse fall within the interpulse interval—for example, the interpulse intervals of QPS are 1.5–100 ms, those of theta burst stimulation are 20 ms, and those of paired pulses range from a few ms to hundreds of ms. The delay may have to be considered for applications where sub-millisecond timing is critical, for example phase-specific TMS for a certain electroencephalogram (EEG) rhythm. Still, the delay could be accounted for by prediction algorithms for the EEG phase [67] and triggering the optimized TMS pulse earlier so that the E-field peak aligns with that of a monophasic pulse.

The second optimization step adjusted the amplitude of the waveform based on stimulation thresholds obtained from simulations with a range of model neurons. The linear membrane model required substantially higher amplitude adjustments than the nonlinear models since the linear model did not capture the reduction of threshold by hyperpolarizing leading phase due to deinactivation of the sodium channels [68–72]. The threshold-based scaling factors were consistent across nonlinear models and cell types, demonstrating robustness of the amplitude adjustment step. Thus, the leading phase has two-fold advantage: it reduces the peak coil current of the main phase and reduces its activation threshold. Importantly, the threshold orientation maps of the optimized waveforms were scaled versions of the maps for monophasic pulses [47] given the small variability (< 10%) of the scaling factors across E-field orientations for the individual Blue Brain model clones, indicating that the directional selectivity was not changed compared to the original pulse.

The optimized waveforms with the least losses produced only about one quarter of the coil heating of conventional monophasic pulses and up to 50% less heating than biphasic pulses. Using these optimized monophasic-equivalent pulses, the pulse rate of TMS protocols can be at least quadrupled or doubled with equal power loss and coil heating compared to conventional monophasic and biphasic pulses, respectively. Further increases in pulse rate can be obtained, considering that the MPS-TMS device recovers pulse energy as in conventional biphasic stimulators [38]. The higher pulse rate can be achieved with reduced interpulse or interburst intervals, increased number of pulses per burst, or a combination of both.

### 4.2. Comparison with other optimization approaches

Waveform optimization approaches have been developed for neural stimulation, and the pulse energy is a common optimization target. Classic optimization methods, such as function parametrization with subsequent gradient decent [73] or calculus of variations and the related principle of least action [74–76], have been applied only to single-or two-compartment neuron and straight axon models. In contrast, metaheuristic methods such as evolutionary algorithms [77–79] and particle swarm optimization [37,63,80,81] can not only deal with the nonlinearity of neural response, but also handle complex neuronal structures, multiple stimulation targets, and activation of neural networks. These methods, however, are computationally expensive as they need to evaluate a large population of candidate solutions over hundreds to thousands of iterations and are sometimes repeated with tens to hundreds of trials to ensure convergence [77,78,81].

Instead of imposing both the E-field matching and neural activation constraints simultaneously during the optimization, our approach used the relationship between TMS coil current and E-field waveform to separate the neural response from the optimization process and therefore reduced the complexity of the problem significantly. The actual optimization had a short computational runtime as the neural simulations were only run once per parameter combination to obtain the scaling factors for adjusting the waveform amplitudes. A single CPU can easily handle the optimization and simulation using simple neural models. Parallel computing on a cluster was used to map the thresholds for all 200 E-field orientations of the 15 Blue Brain models within practical runtime, or similarly to find the threshold distribution of a population of 3000 neurons in the motor cortex of a realistic head model [47]. For example, distributing the 200 simulations over 50 CPUs reduced the runtime for the Blue Brain models to approximately 10 minutes (L2/3 pyramidal neurons) to 2 hours (L4 interneuron, clone 2) for a given combination of parameters. Nevertheless, the computational cost was significantly less compared to metaheuristic methods, which typically take many days to complete even on a compute cluster.

Previous efforts to optimize TMS waveforms minimized coil heating without constraining the pulse shape to any equivalency while simultaneously accounting for neural activation threshold [37,63]. By only constraining the coil voltage, these optimizations generated waveforms that had a negative leading phase and a triangular main phase for the current and rectangular biphasic shapes for the E-field (Figure 4B). With the negative leading phase and the short main phase, these optimized waveforms from previous work generated less heating compared to all three types of conventional pulses. However, their stimulation characteristics would resemble those of a half-sine pulse, which tend to have different thresholds, activation latencies, and directional selectivity than monophasic pulses [9,11,47].

In comparison, the optimized waveforms in the present work achieved similar performance in terms of heating compared to two of the three previous waveforms (blue and red data points in Figure 4A, corresponding to coil voltage constraints of 990 V and 1960 V in [37], respectively), while retaining the excitation characteristics of conventional monophasic pulses.

### 4.3. Limitations

Our optimization method used the time derivative relationship between the TMS coil current and E-field to generate pulses with equivalent neural activation properties compared to an E-field waveform template. Therefore, the performance of the method depends on the specific pulse template and could not be generalized. For example, shifting conventional half-sine and biphasic waveforms in ways similar to the shift in the optimized waveforms also reduced coil heating, albeit to a lesser extent, and after the threshold-based amplitude adjustment, the reduction in heating of the optimized half-sine and biphasic waveforms was further diminished (Figures S12 and S13). Whereas a 8%−55% reduction in heating was achieved for half-sine waveforms (based on results of nonlinear models), the biphasic waveforms already have an average coil current close to zero and therefore the heating was largely unchanged (within 10%) by the optimization. Accordingly, our approach is most effective for mostly unidirectional current pulses, e.g., monophasic and half-sine pulses, whereas other methods are needed to design energy-efficient waveforms with excitation properties equivalent to other pulse shapes.

## 5. Conclusion

We developed a two-step waveform optimization method to design TMS pulses that produced significantly lower coil heating compared to conventional monophasic pulses but retained stimulation effects equivalent to the latter. The first optimization step exploited the temporal relationship of the coil current and the induced E-field to generate candidate waveforms without the necessity of performing neural simulations. The final optimized waveforms were obtained by adjusting the amplitude of the candidate waveforms to account for differences in neural activation thresholds. The optimized pulses dissipated as little as 25% of the energy loss of a standard monophasic pulse waveform. Such aggressive reductions of coil heating and power losses in the device, in combination with modern device technology that recovers pulse energy, can enable rapid-rate stimulation protocols while maintaining the orientation selectivity and changes in excitability produced by monophasic pulses.

## Acknowledgments

This work was supported by Grant No. RF1MH124943 from the National Institute of Mental Health of the National Institutes of Health of the United States of America; the content is solely the responsibility of the authors and does not necessarily represent the official views of the funding agency. Computational support was provided by the Duke Compute Cluster.

## Availability statement

The code and data that support the findings of this study are available online at the Research Data Repository (RDR) of Duke University Libraries: https://doi.org/10.7924/r4wd41f7d.

## Author contribution

S. M. Goetz, A. V. Peterchev, and W. M. Grill conceived, supervised, and obtained funding and computational resources for the study. B. Wang performed the waveform optimization and neuron modeling, data analysis, and visualization. Z. Li and J. Zhang developed the MPS-TMS system and control program. J. Zhang performed the experiment and recorded the optimized waveforms. B. Wang, J. Zhang, and S. M. Goetz wrote the manuscript, and all authors revised, commented on, and approved the final version of the manuscript.

## Supplement

**Figure S1:**
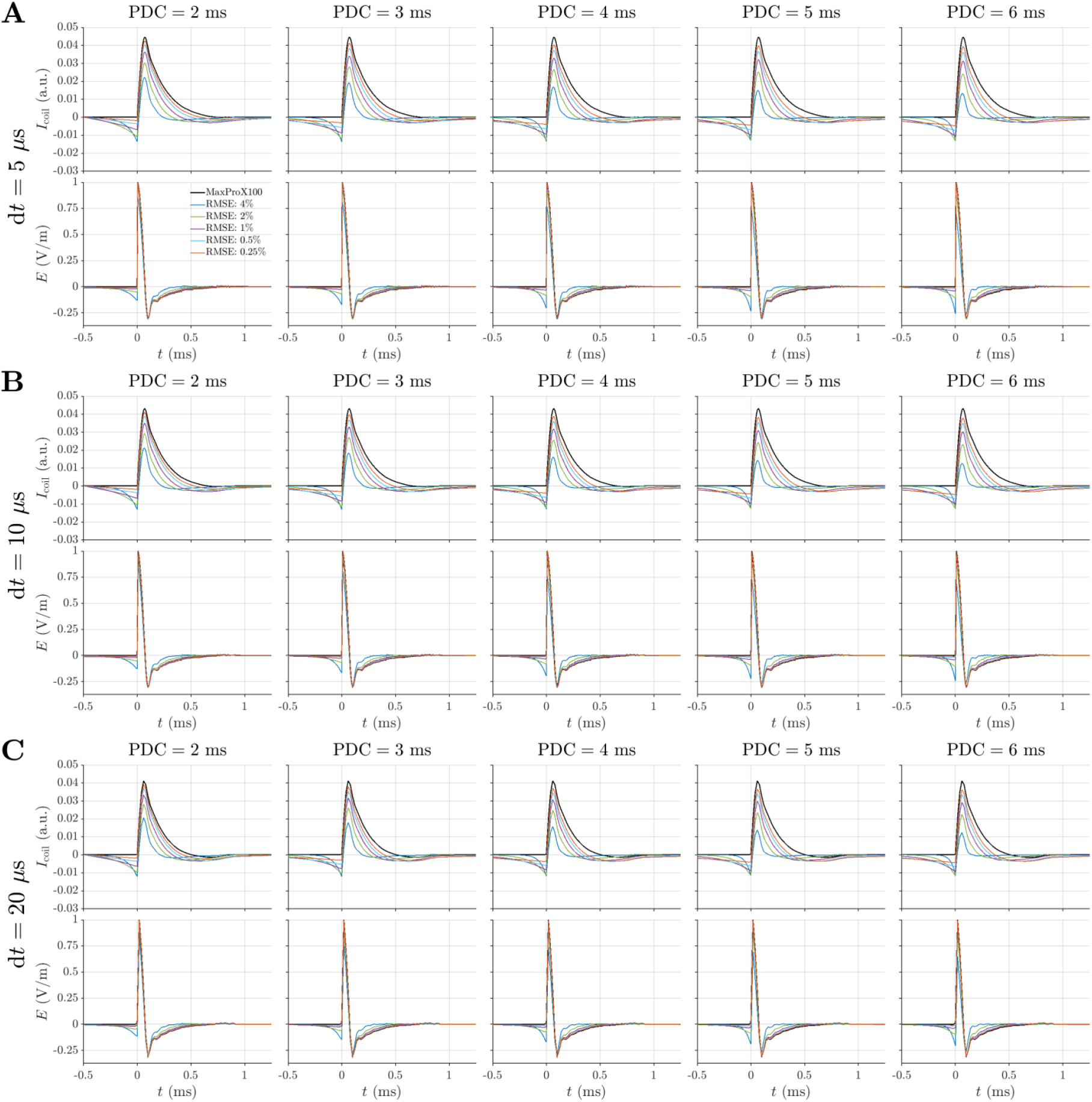
Waveform optimization with different time steps. Similar to Figure 2A, with results of all RMSE and pulse duration constraints for **A** 5 μs, **B** 10 μs, and **C** 20 μs optimization time steps, respectively. Besides the difference in waveform smoothness for the different time steps, the coil currents have slight amplitude difference due to the numerical accuracy of the integration.

**Figure S2:**
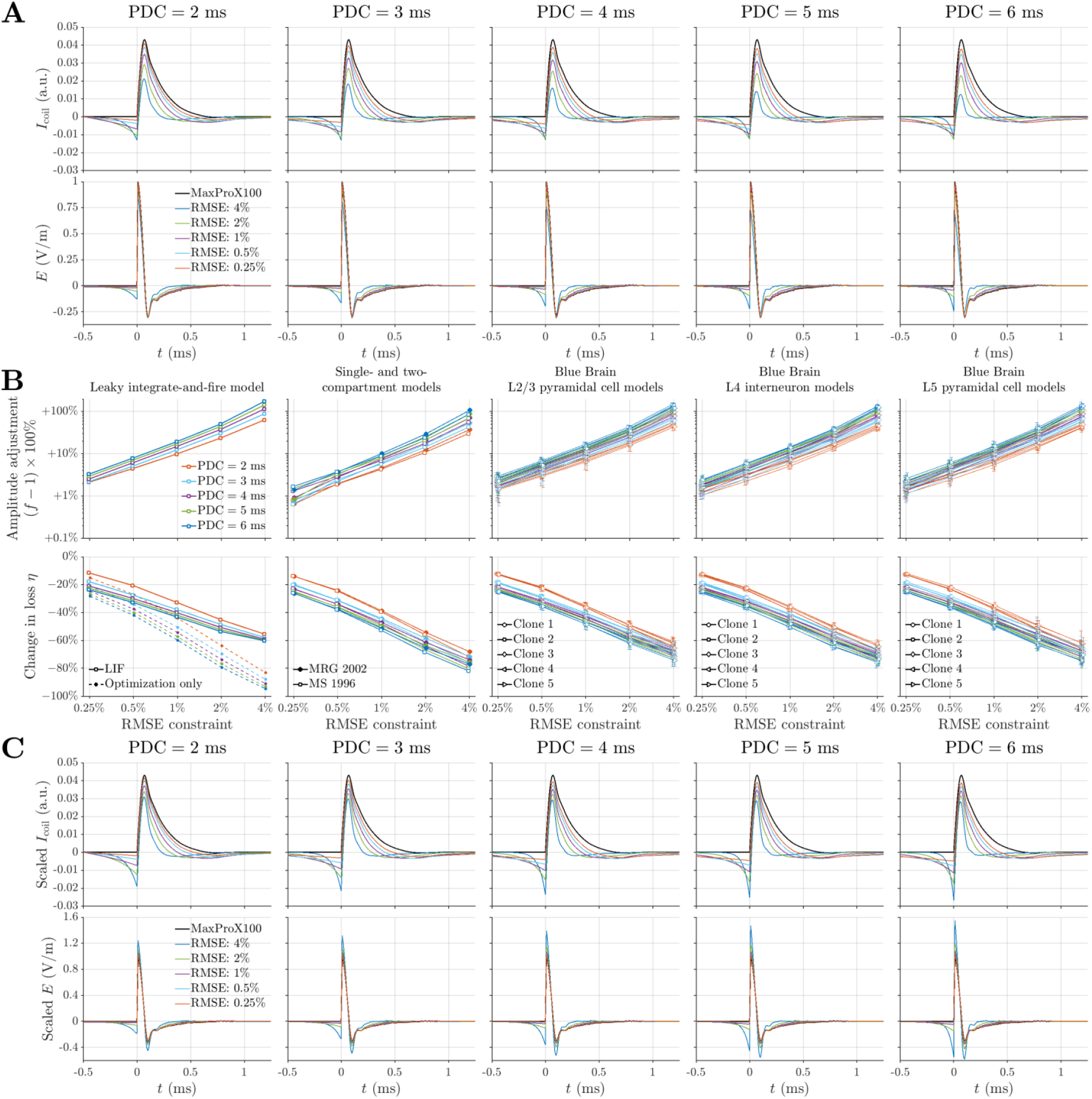
Amplitude adjustment and optimized waveforms for monophasic pulses. Similar to Figures 2 and 3, with results for all RMSE and pulse duration constraints and model clones. Error bars in B show standard deviation across 200 E-field orientations for each clone. Only computed waveforms are shown in C, without the scaling to match measurement.

**Figure S3:**
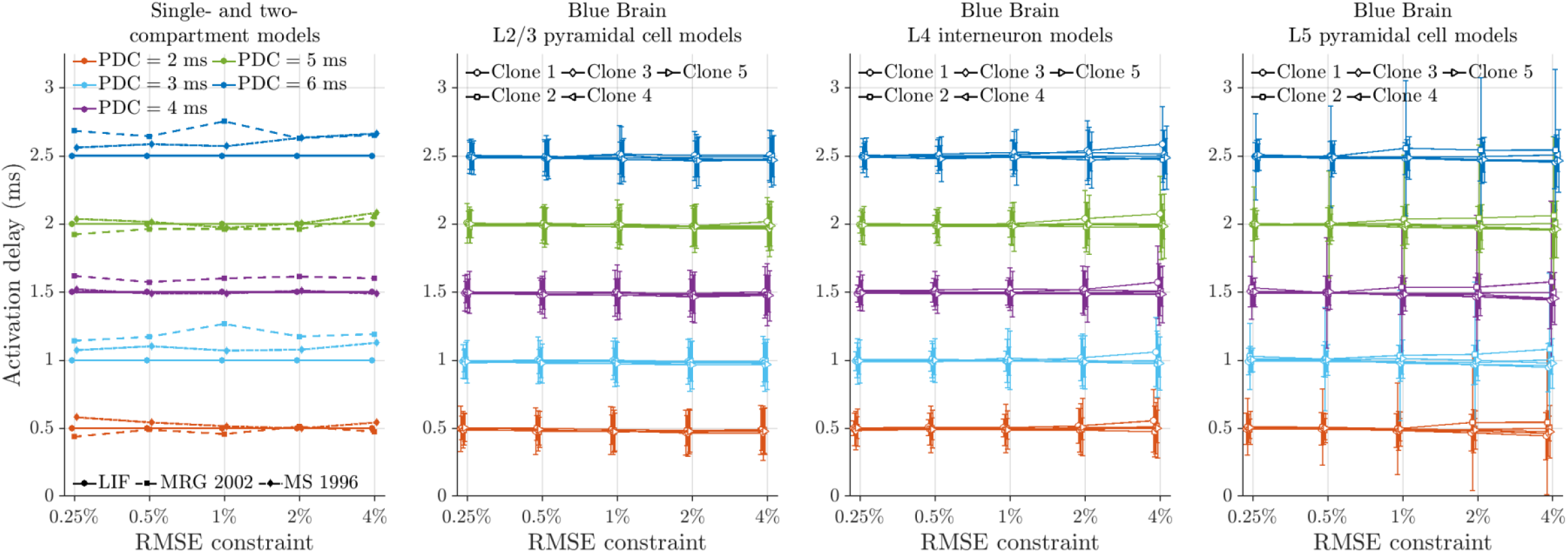
Delay of neural activation. The delay of the action potentials generated by the optimized pulses, including the leading phase, compared to the standard monophasic pulse. Considering the main phase only, the delay was zero with small variability. Errors bars show mean and standard deviation of 200 E-field orientations for the Blue Brain models. PDC: pulse duration constraint. LIF: leaky integrate-and-fire model MRG: McIntyre–Richardson–Grill model. MS: Mainen–Sejnowski model.

**Figure S4:**
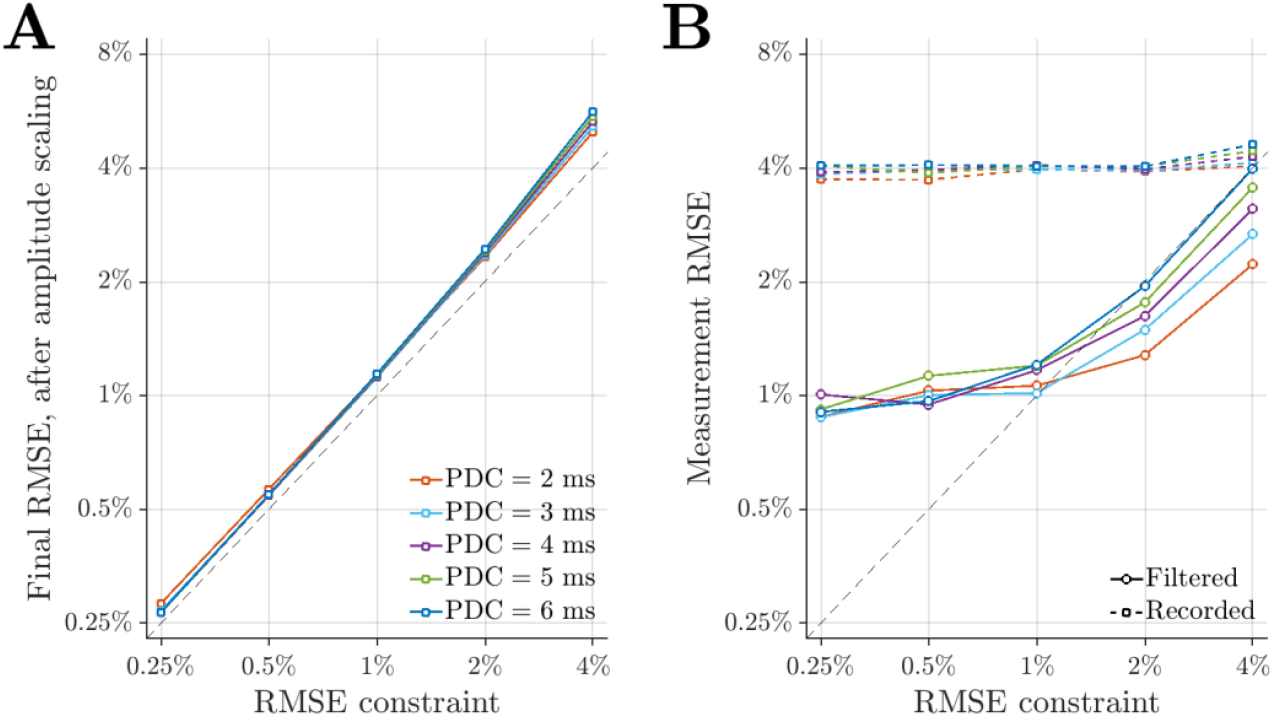
RMSE of computed and measured optimized waveforms. RMSE of the final optimized waveform versus the original TMS waveform for computed results (**A)** and recorded waveforms (**B**). The amplitude adjustment step increased the RMSE of the optimized waveforms compared to the RMSE constraint in the first optimization step. The RMSE of the unfiltered waveforms were subject to noise and around 4% for any optimization parameter, whereas the filtered waveform had either larger or slightly smaller RMSE compared to the RMSE constraint due to the distortion of the filtering process. PDC: pulse duration constraint.

**Figure S5:**
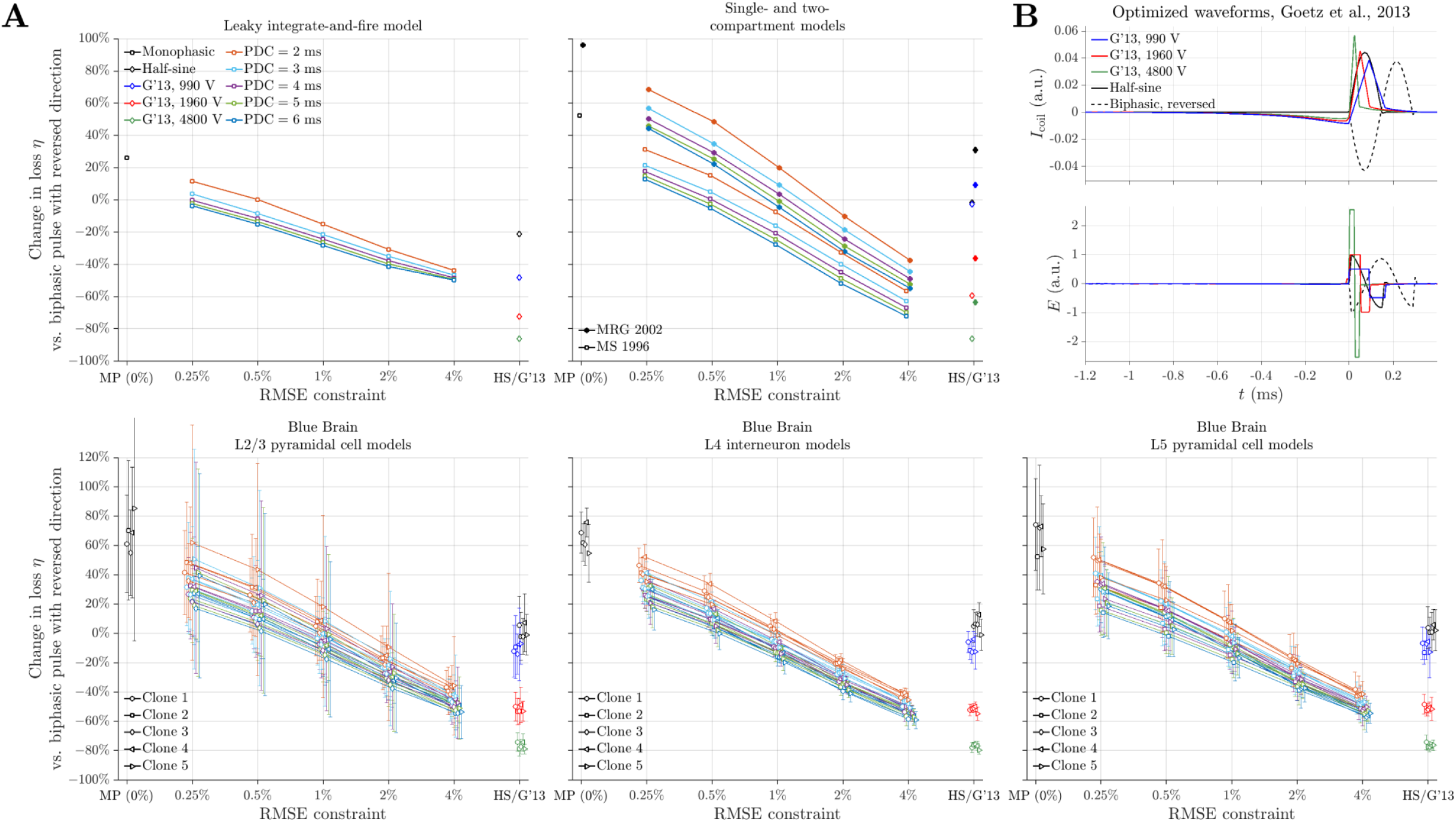
Difference in coil heating comparing monophasic-like pulses and optimized waveforms from a previous study against biphasic pulse. Similar to Figure 4, with results for all pulse durations and Blue Brain model clones in panel A and the full waveform shown for the optimized waveforms from previous work in panel B.

**Figure S6:**
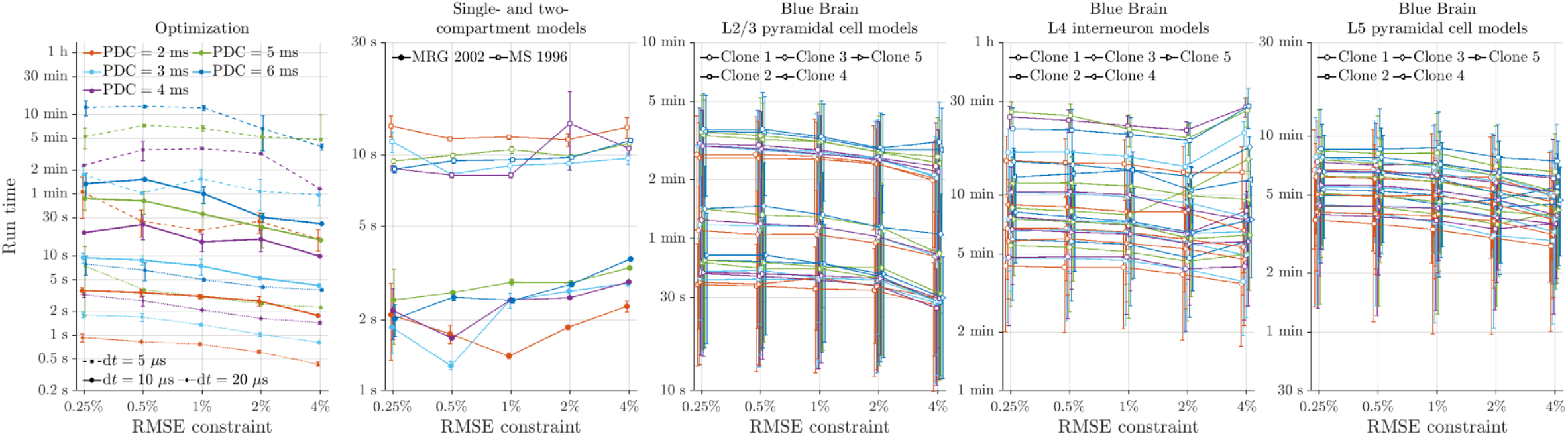
Runtime of optimization and neural simulations. The runtime of the waveform optimization (mean and standard deviation of ten trials for 10 μs and 20 μs time steps, and four trials for 5 μs time step) and threshold search for the single- and two-compartment models (mean and standard deviation of ten trials), and the Blue Brain L2/3 pyramidal cell, L4 interneuron models, and L5 pyramidal cell models (mean and standard deviation of 200 E-field orientations). The runtime of the leaky integrate-and-fire model was on the order of milliseconds and not shown. PDC: pulse duration constraint. MRG: McIntyre–Richardson–Grill model. MS: Mainen–Sejnowski model.

**Figure S7:**
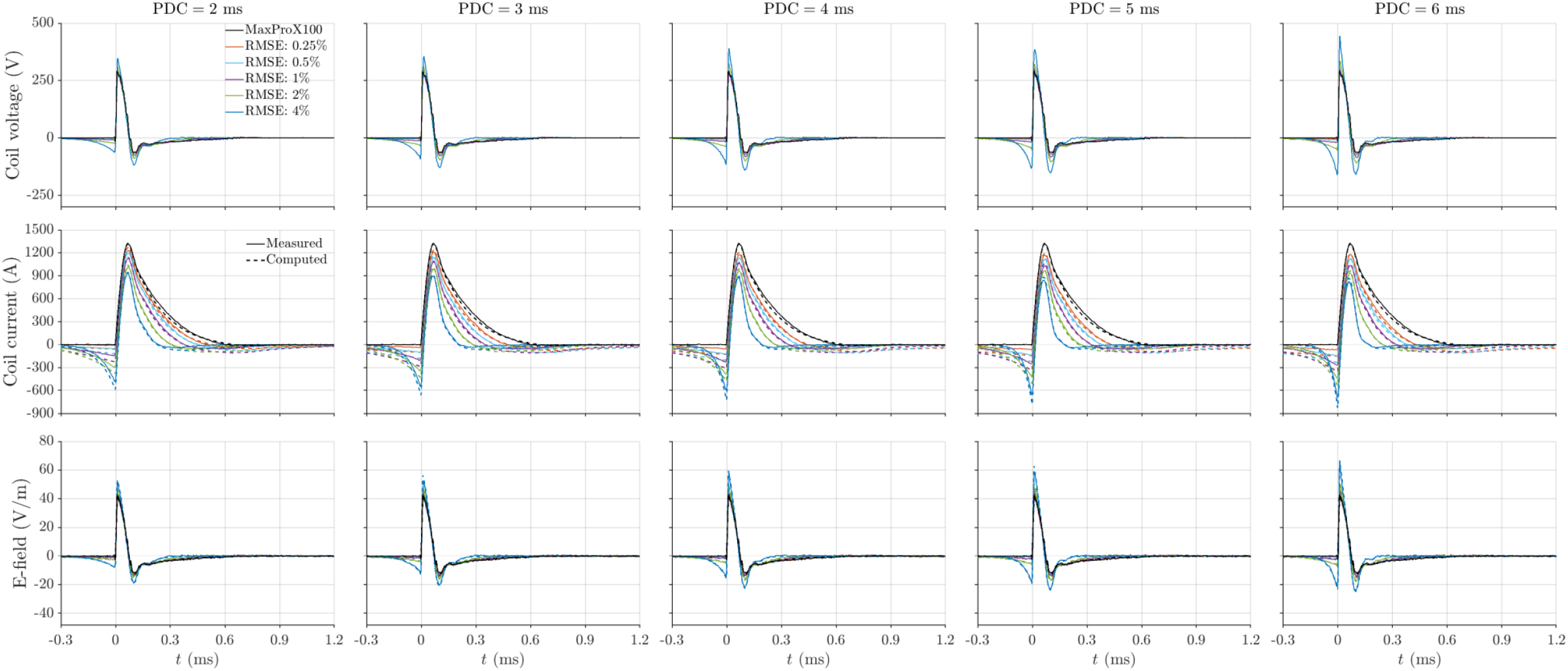
Computed and measured optimized TMS waveforms. Similar to Figure 3, with results for all RMSE and pulse duration constraints.

**Figure S8:**
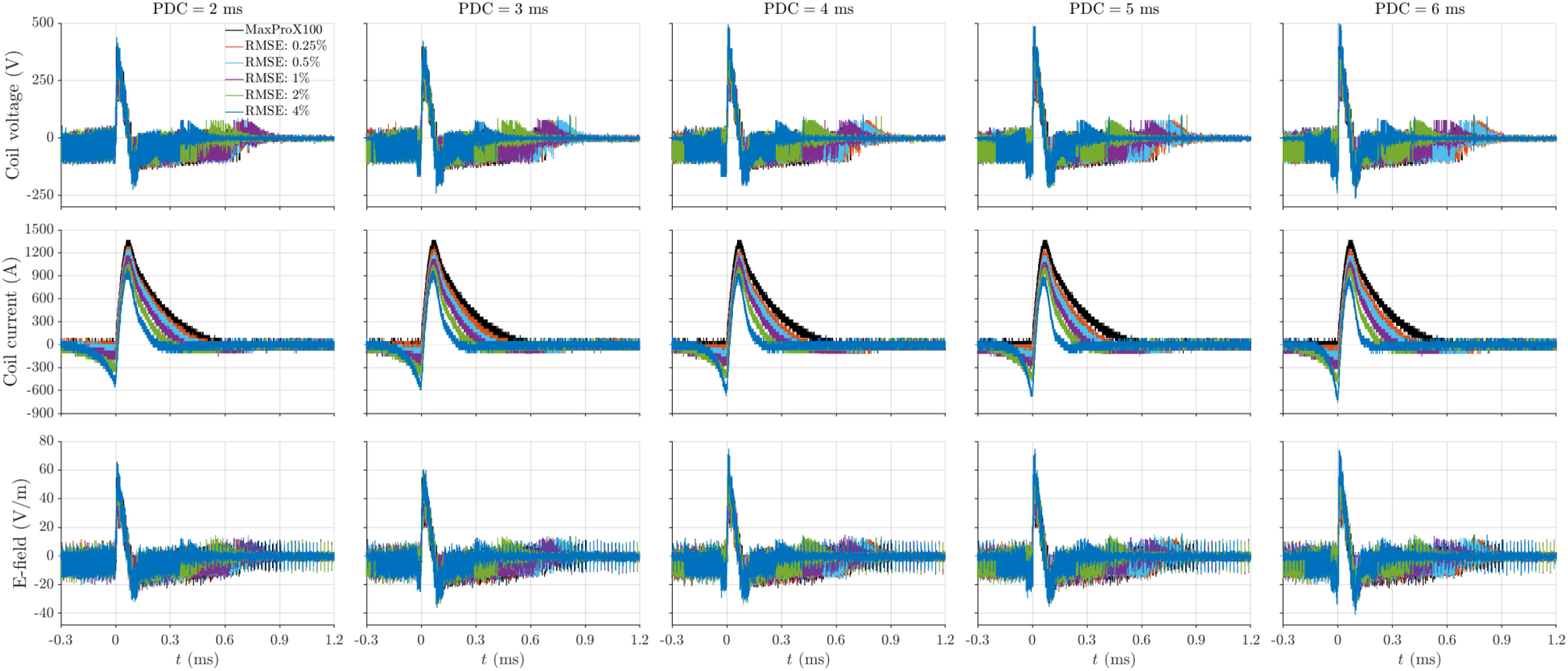
Measured optimized TMS waveforms. Similar to Figure S7, with unfiltered measurements and without the computed waveforms.

**Figure S9:**
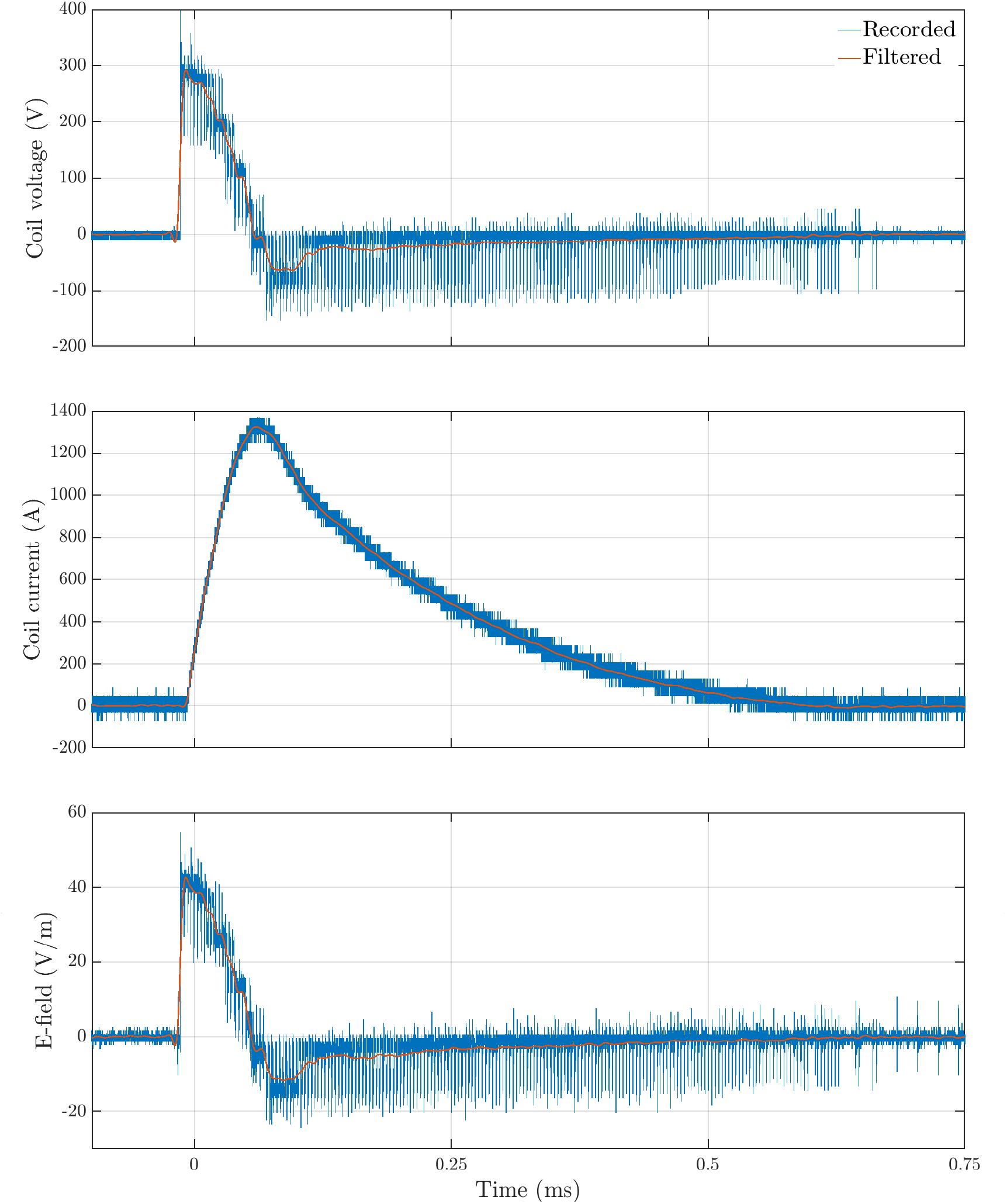
Raw and filtered recordings of the MaxProX100 TMS pulse implemented by MPS-TMS. The coil current contains mostly high-frequency measurement and quantization noise, whereas the coil voltage and E-field waveforms also include switching noise. Filtering removes most of the noise, but the voltage levels of the multilevel waveform synthesis are still visible.

**Figure S10:**
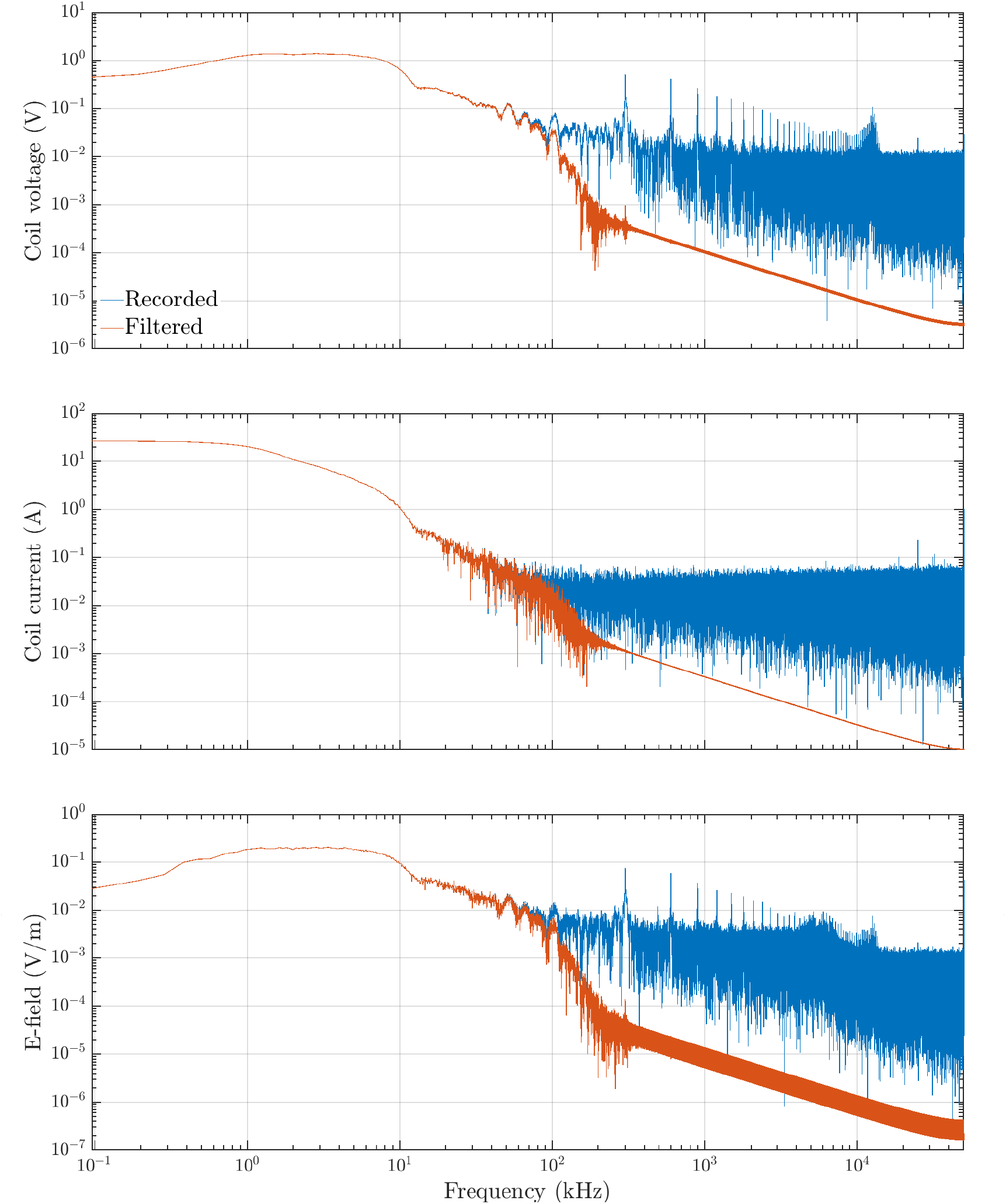
Spectra of raw and filtered recordings of the MPS-TMS-implemented MaxProX100 TMS pulse. The spectra contained noise in the high frequency range above 100 kHz, which is significantly reduced by filtering. The transistors’ switching noise with fundamental frequency of 300 kHz is visible in the spectra of the coil voltage and E-field.

**Figure S11:**
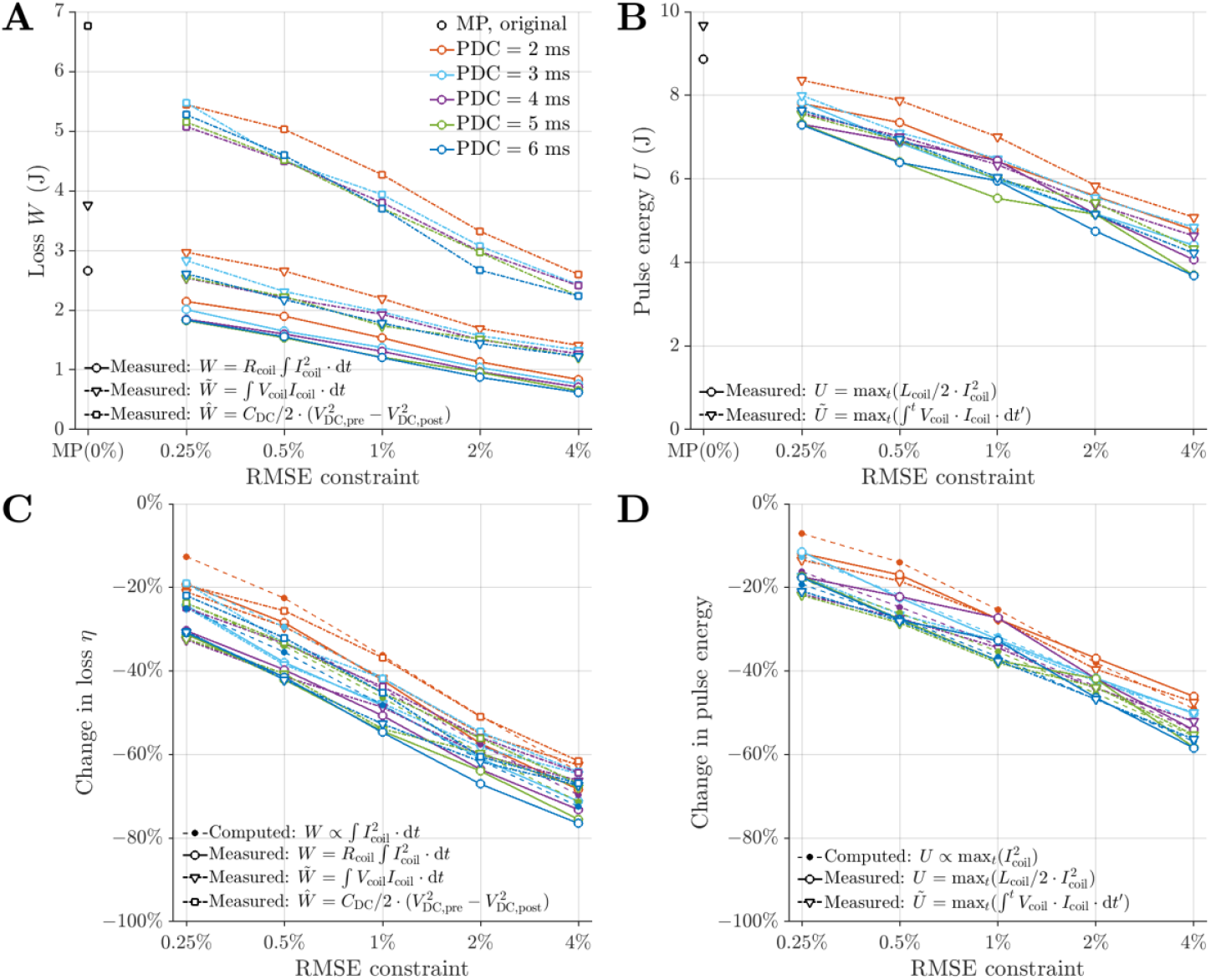
Losses and pulse energy of optimized pulses and original TMS pulse. Similar to Figure 5, with results for all pulse duration constraints.

**Figure S12:**
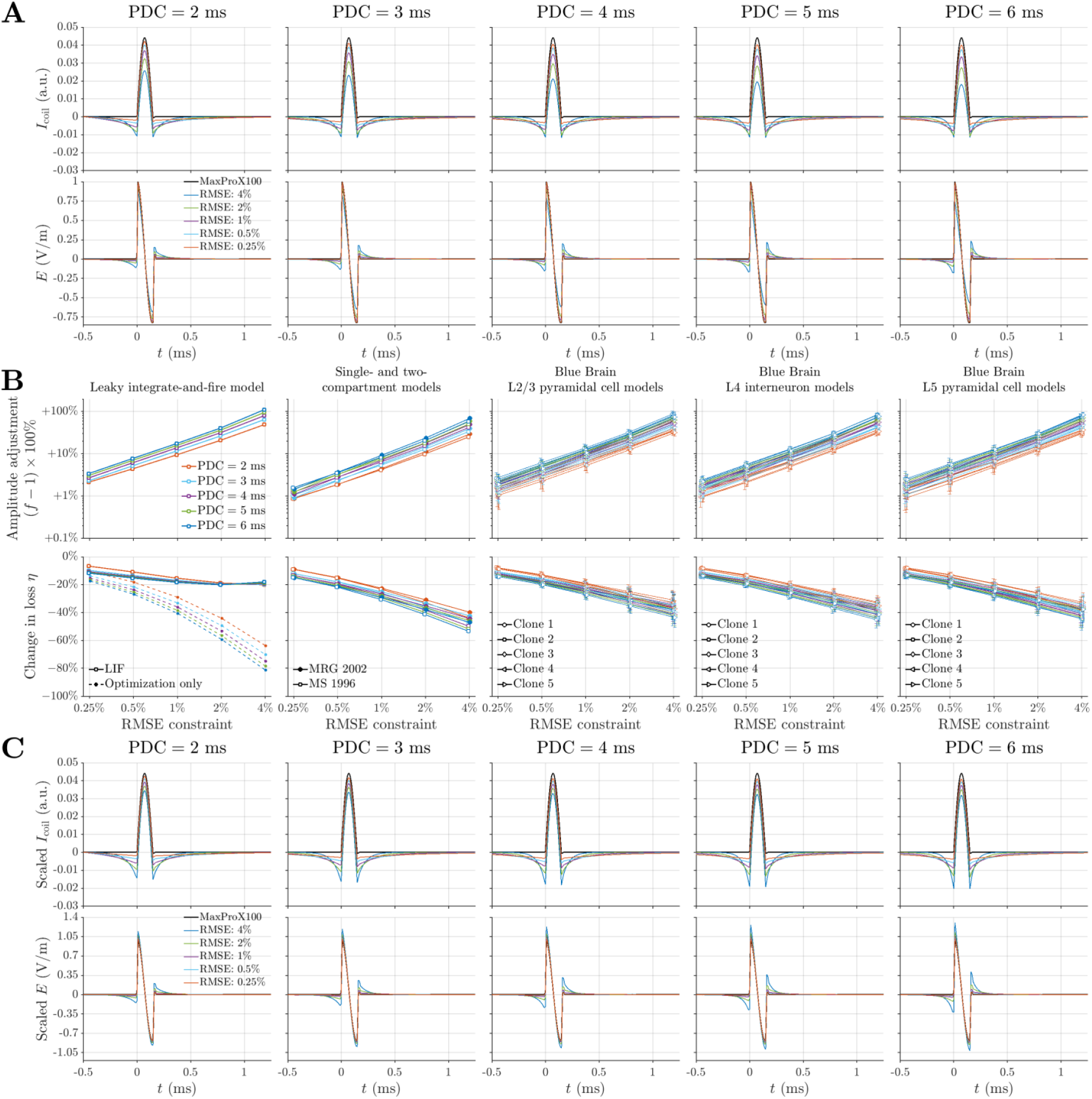
Waveform optimization, amplitude adjustments, and optimized waveforms for half-sine TMS pulses. Similar to Figure S2.

**Figure S13:**
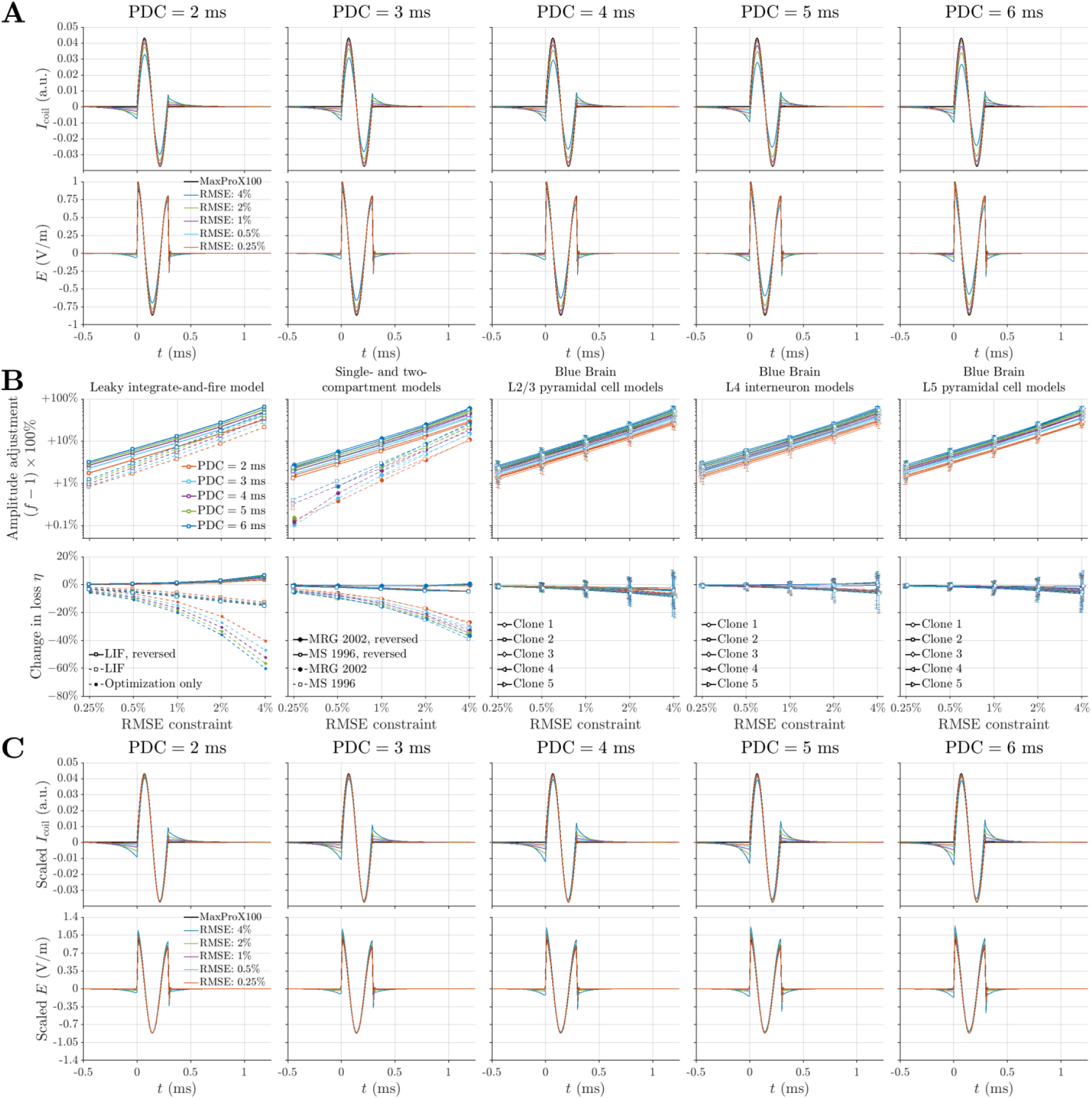
Waveform optimization, amplitude adjustments, and optimized waveforms for biphasic TMS pulses. Similar to Figure S2. Amplitude adjustment and change in energy loss for single- and two-compartment models show results for both normal and reversed directions of the stimulation waveform in panel B, with results of the reversed waveform in agreement with those of the Blue Brain models. The Blue Brain models sampled the distribution of E-field orientations, therefore stimulation was effectively applied with the reversed waveforms for each orientation when the normal direction was used for the opposite orientation. Results of single- and two-compartment models using the normal direction showed considerable more reduction of coil heating, however, such achievement were not relevant given the higher threshold and loss compared to those for reversed direction.

